## Supplemental Figures for "Eye movements reveal spatiotemporal dynamics of visually-informed planning in navigation"

### Supplemental materials

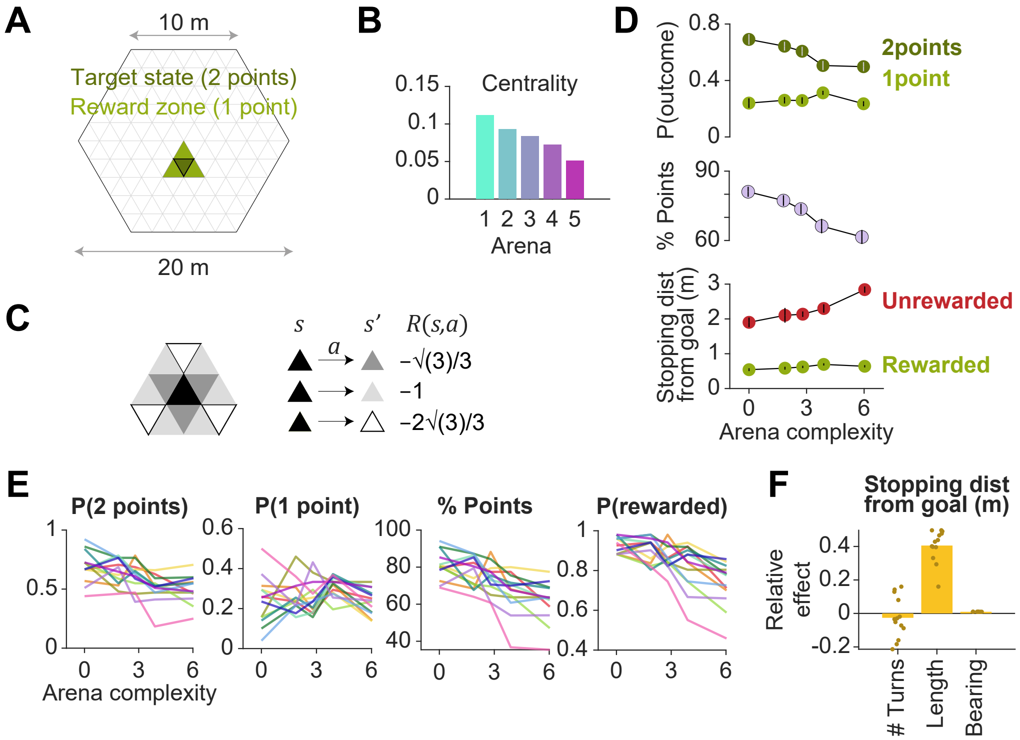


*Figure S1:* **A.** Arenas were regular hexagons of side length 10 m with a triangular tessellation of side length 2 m. Two points were rewarded if participants reached the goal state (green), and one point was rewarded if participants reached a state neighboring the goal state (light green). **B.** Mean state closeness centrality of each arena, where higher centrality values correspond to a less complex arena. Error bars denote $\pm 1$ SD across states. **C.** To incorporate twelve degrees of freedom in translation, value functions were computed using dynamic programming, whereby the cost of actions scaled in accordance with the center-to-center distance between states $s$ and $s'$ (pertaining to the transition which results from taking action $a$). **D.** Top: Across all participants and all trials, the probability of being awarded two points (green) decreased with arena complexity, while the probability of being awarded one point (light green) was relatively constant across all arenas. Middle: The total fraction of points earned decreased with arena complexity. Bottom: Distance between the stopping location and the goal in rewarded (green) and unrewarded (red) trials. Error bars denote $\pm 1$ SEM. **E.** Each color corresponds to a single participant. Variables plotted against arena complexity are, from left to right: the probability that participants scored two points or one point, the percentage of total points scored, and the probability of being rewarded. **F.** Linear mixed model with random intercepts and slopes for the effect of trial-specific variables (number of turns, length of optimal trajectory, relative bearing, and the number of trajectory options) on the distance between the participant's stopping position and the goal (error). The path length best predicts stopping errors. The overlaid scatter shows fixed effect slope + participant-specific random effect slope, and all variables were z-scored prior to model fitting.

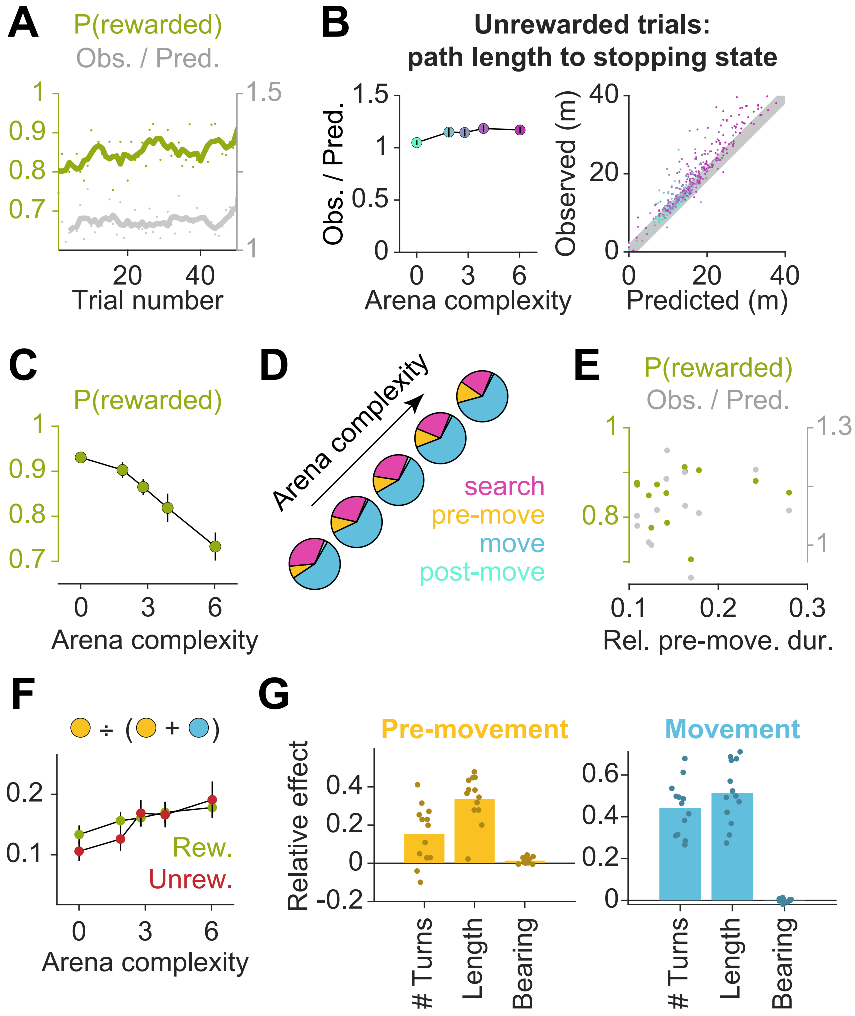


*Figure S2:* **A.** Performance was stable across each block, as measured by the average probability of being rewarded on each trial (green), as well as the average ratio between the empirical and optimal path lengths (gray). **B.** Left: Across all arenas (colored according to the coloring scheme introduced in Figure 1b), the path lengths observed in unrewarded trials were close to the optimal trajectory lengths between the starting state and the state at which participants stopped on these trials, suggesting that unrewarded trials were predominantly caused by participants forgetting the precise location of the target. Right: The ratio of observed to optimal path lengths (to the participants' stopping location on unrewarded trials) was close to unity in all arenas. **C.** Arena complexity predicts the fraction of rewarded trials in each arena. **D.** Distribution of epoch durations across all participants and all trials. Pre-movement and movement occupied a greater fraction of the total trial time for less open arenas. **E.** Some participants spent less time deliberating before movement, but this did not impact task performance. Relative pre-movement duration was defined as the average ratio of the duration of the pre-movement epoch to the duration of the entire trial after goal detection. The average proportion of time that participants spent making prospective eye movements prior to using the joystick did not correlate with their average path lengths across all arenas (gray), nor with the overall probability of them being rewarded (green). **F.** The relative pre-movement duration did not differ between rewarded (green) and unrewarded trials (red), except for the two easiest arenas (where planning demands are low). This suggests that unrewarded trials are not merely caused by poor planning. **G.** Linear mixed models with random intercepts and slopes for the effect of trial-specific variables on epoch durations – left: pre-movement; right: movement; and relative pre-movement duration, top right. (Please see the description for Figure S1f for further details.) The number of turns and the length of the optimal trajectory had the greatest effect on epoch durations. All error bars denote $\pm 1$ SEM.

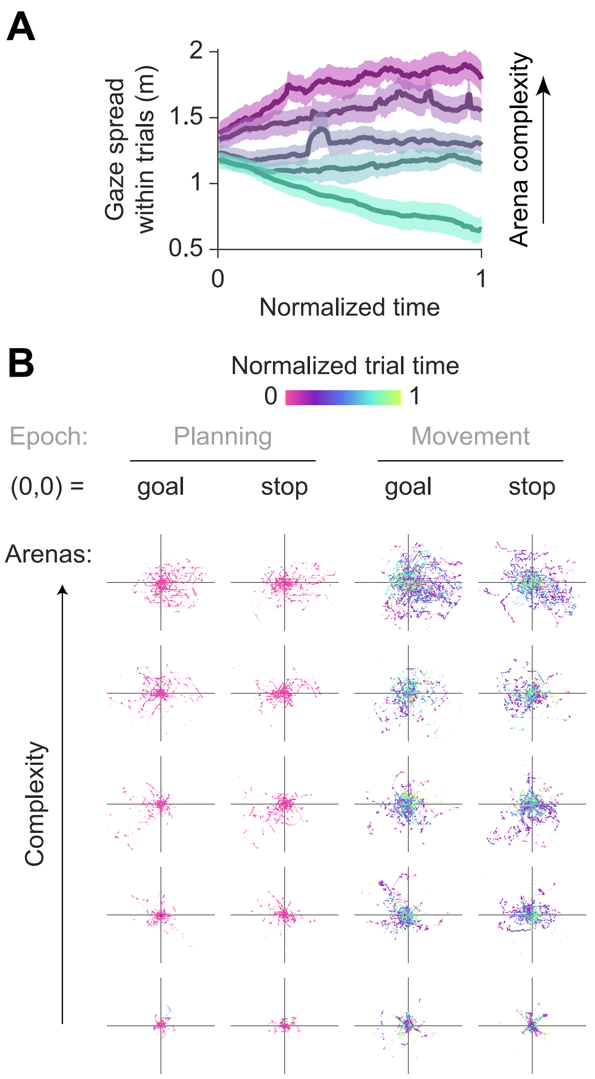


*Figure S3:* **A.** The variance of gaze within trials across reaches a peak in the second half of the pre-movement period. Because the pre-movement duration scales with arena complexity, variance was computed using a sliding window of length between 0.32 s (least complex arena) to 0.68 s (most complex arena), linearly spaced between the extremes for arenas of intermediate complexity. Endpoints that fell outside the window were discarded. Results were qualitatively similar when using a fixed window size for all arenas. **B.**Gaze is increasingly concentrated around the goal/stopping location for easier arenas. The believed goal location was assumed to be the participants' stopping position, and the point of gaze was visibly more concentrated around the stop location than around the true goal location (especially in the most complex arena). The effects of working memory on gaze were more apparent for easier, more open arenas. Each panel depicts eye movements on a random subset of trials (3 trials x 13 participants) in each arena. The origin (0,0) denotes the goal location or the stopping location. Raw gaze positions relative to these points are depicted during the pre-movement and movement epochs. Axis limits are $\pm$15 m for all panels.

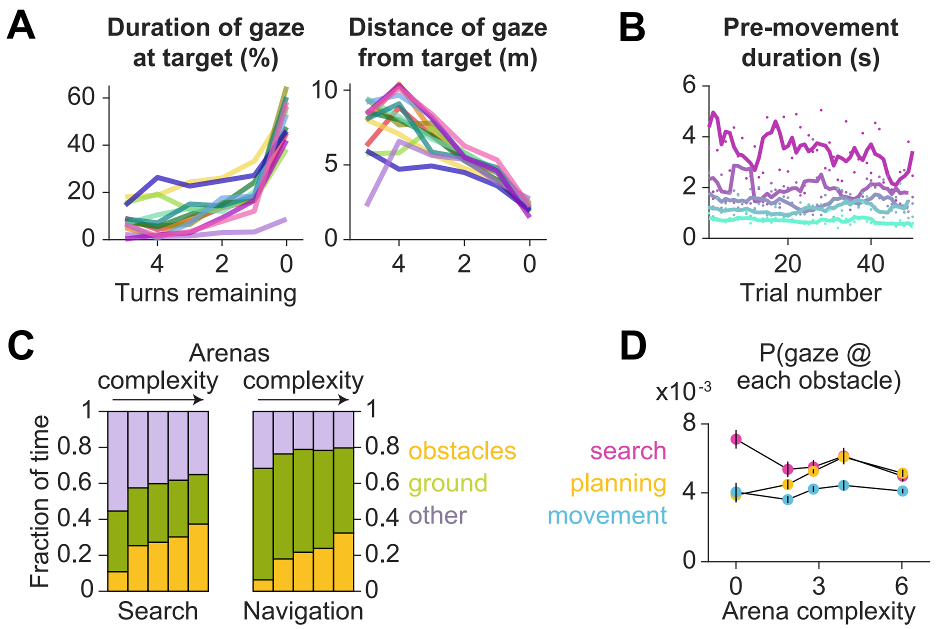


*Figure S4:* **A.** Each color corresponds to a single participant. Variables plotted against arena complexity are — left: percent of time participants spent gazing at the target location, right: average distance of gaze to the target location. **B.** The pre-movement duration in each arena (colored according to the color scheme used for Figures 1b and S1b) does not significantly change across trials. **C.** During search, participants spent a greater fraction of time foveating the arena borders (purple), and after the target was located, participants spent more time foveating the ground (green). While there appears to be a trend in the fraction of time foveating obstacles (orange) vs. arena complexity, this is explained by a higher obstacle density in the more complex arenas. **D.** Across all participants and all trials, the probability of gazing upon each obstacle remains relatively constant across all arenas during each epoch. All error bars denote $\pm 1$ SEM across participants.

**Relevance simulations:** Motivating the quantitative characterization of the task relevance of visual samples, we show an illustration of the consequence of mistaken beliefs about the passability of specific transitions on the subjective value function. Transition ***toggling*** can be defined as the act of removing an obstacle between two states if an obstacle was previously present, or adding an obstacle at that location if one was previously absent. In alignment with intuition, some transitions are more important to veridically represent (Figure S5a – middle), as toggling them results in a dramatic change to the value function (which is essential to computing the optimal set of actions to reach the goal), while toggling some other transitions causes a relatively minimal change (Figure S5a – right).

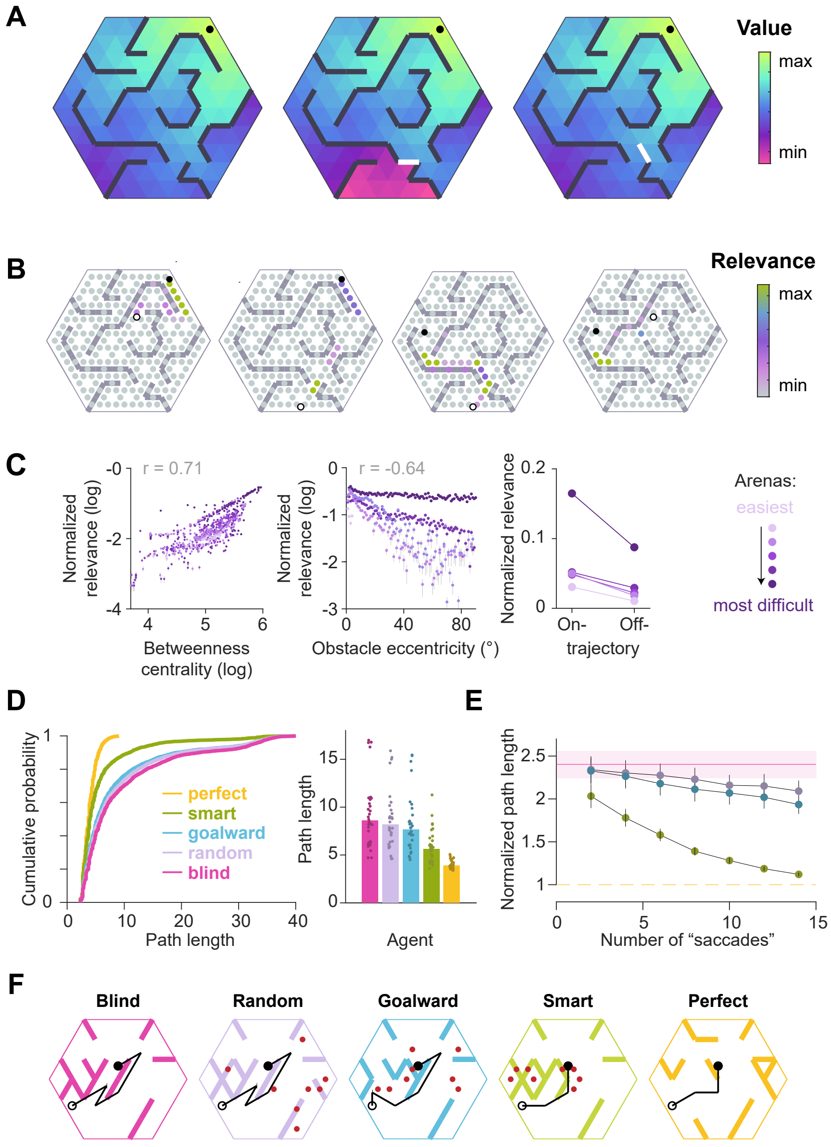

*Figure S5:* **Simulations validate the utility of precisely knowing the status of theoretically important transitions. A.** Value functions corresponding to an arbitrary goal location (closed circle) in an example arena (left) and in the arenas resulting from blocking either a bottleneck transition (center) or a transition that was not a bottleneck (right). **B.** Theoretical relevance of all transitions (circles) for the example arena for four different pairs of start (open black circle) and goal (closed black circle) states. **C.** The betweenness centrality of a state describes the degree to which the state controls the traffic flowing through the area (see Methods). Left: Across all possible start and end locations, the mean normalized relevance of non-obstacle transitions (across all possible start and goal state pairings) was positively correlated with betweenness centrality values. Middle: The mean normalized relevance of obstacles was negatively correlated with the eccentricity of each obstacle from the straight line connecting the current state to the goal state. Right: Transitions that fell on the optimal trajectory had greater relevance than those that fell outside of it. **D.** Simulation results of an agent instantiated with a perfect transition model (orange), and four agents with imperfect transition models, three of whom were endowed with the ability to correct their model according to different rules (see text). Those three agents were allowed to make eight 'saccades', each of which could update one transition. Left: Cumulative distributions (CDFs) of the path lengths of various agents (100 trials each from 25 different arenas; see Methods). Right: Median of trial-averaged path lengths across all simulated arenas; data points denote trial-averaged path lengths in individual arenas. **E.** Results of simulations similar to **D** but with a variable budget of 'saccades'. Each line denotes the average path length (across arenas) of one agent as a function of the number of 'saccades'. For each trial, path lengths of different agents were normalized by the optimal path length before trial-averaging. Error bars denote $\pm1$ SEM. **F.** Example simulated trajectories, as well as the gaze samples (red dots, if applicable), taken by each agent. The configuration of the arena reflects the agent's subjective model at the *end* of all eye movements. Note that the subjective model of the "smart" agent was still quite mismatched with the true world model after eight eye movements, but the visual samples allowed for the correction of the model at crucial locations such that the trajectory of the "smart" agent was closer to optimal than that of the other agents.

---

By defining relevance of transitions according to Equation 1, we can thus capture multiple task-relevant attributes in a succinct manner. Theoretically investigating whether looking at task-relevant transitions improves navigational efficiency, we simulated artificial agents performing the same task that we imposed upon our human participants. One agent ("perfect") had a veridical subjective model of the environment, and thus was capable of computing the optimal trajectory (Figure S5f). Its antithesis ("blind") had an incorrect subjective model where half of the obstacle positions were 'misremembered' (toggled) to simulate a predicament where the agent had previous exposure to the arena, but were only halfway to learning the precise transition structure. The blind agent was incapable of using vision to correct their model prior to taking actions according to their subjectively computed value functions. Performance at these two extremes was compared against the performance of three agents that were allocated a fixed budget of 'saccades' to rectify their incorrect models. These agents either randomly interrogated transitions ("random"), preferentially sampled transitions along the direction connecting the agent's starting location to the goal location ("goalward"), or chose the most task-relevant transitions as defined by the relevance metric in Eq 1 ("smart").

While all three agents showed an improvement over the "blind" agent, the agent with knowledge about the most task-relevant transitions resulted in much shorter average path lengths than agents looking at transitions along the general direction of the goal or looking at random transitions (mean path length $\pm$ SE – perfect: 3.9$\pm$0.1, blind: 9.6$\pm$0.7, random: 8.9$\pm$0.6, goalward: 8.6$\pm$0.7, smart: 5.8$\pm$0.3; Figure S5d). Moreover, the performance of the smart sampling agent quickly approached optimality as the number of sampled transitions increased (Figure S5e). The rate of performance improvement was substantially slower for the goalward and random samplers (linear rather than exponential). These results were robust to the precise algorithm used to compute the value function in Eq 1. In particular, the successor representation (SR) has been proposed as a computationally efficient, biologically plausible alternative to pure model-based algorithms like value iteration for responding to changing goal locations [3, 27]. We found that estimating the task-relevance of transitions using values implied by SR resulted in a similar performance improvement (Figure S6). Nevertheless, we emphasize that our objective was to use the relevance metric simply as a means to probe whether humans preferentially looked at task-relevant transitions. Understanding how the brain might compute such metrics is outside the scope of this study.

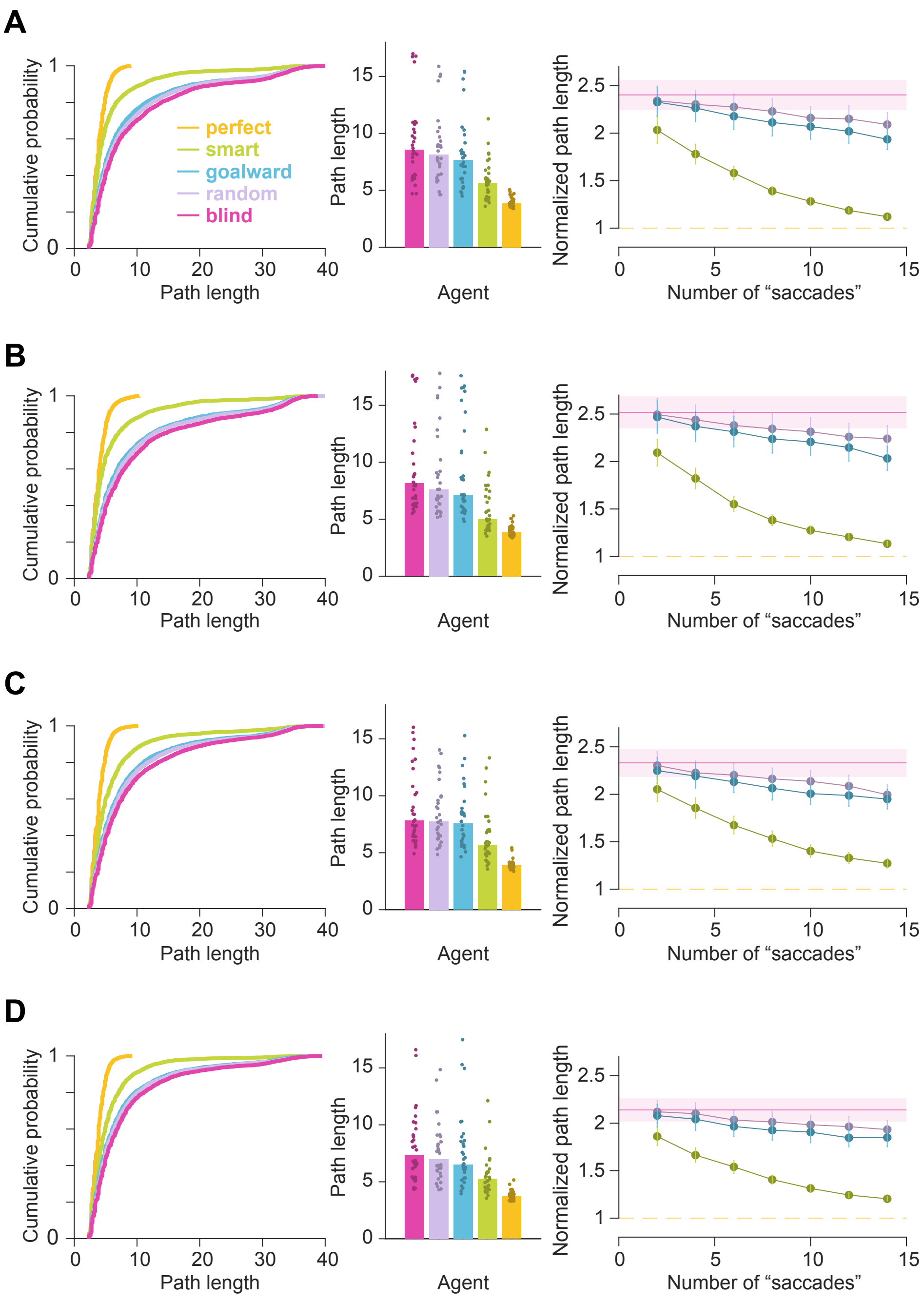


*Figure S6:* Simulations reveal that foveating ‘relevant transitions’ reduces path length. Results were robust to the precise algorithm (value iteration vs. successor representation) as well as the degree of temporal abstraction (current state vs. optimal trajectory) used to estimate the relevance of transitions. Plots similar to Figure 3c and 3d are shown for relevance values calculated with **A** value iteration, current state, **B** value iteration, entire trajectory, **C** successor representation, current state, and **D** successor representation, entire trajectory.

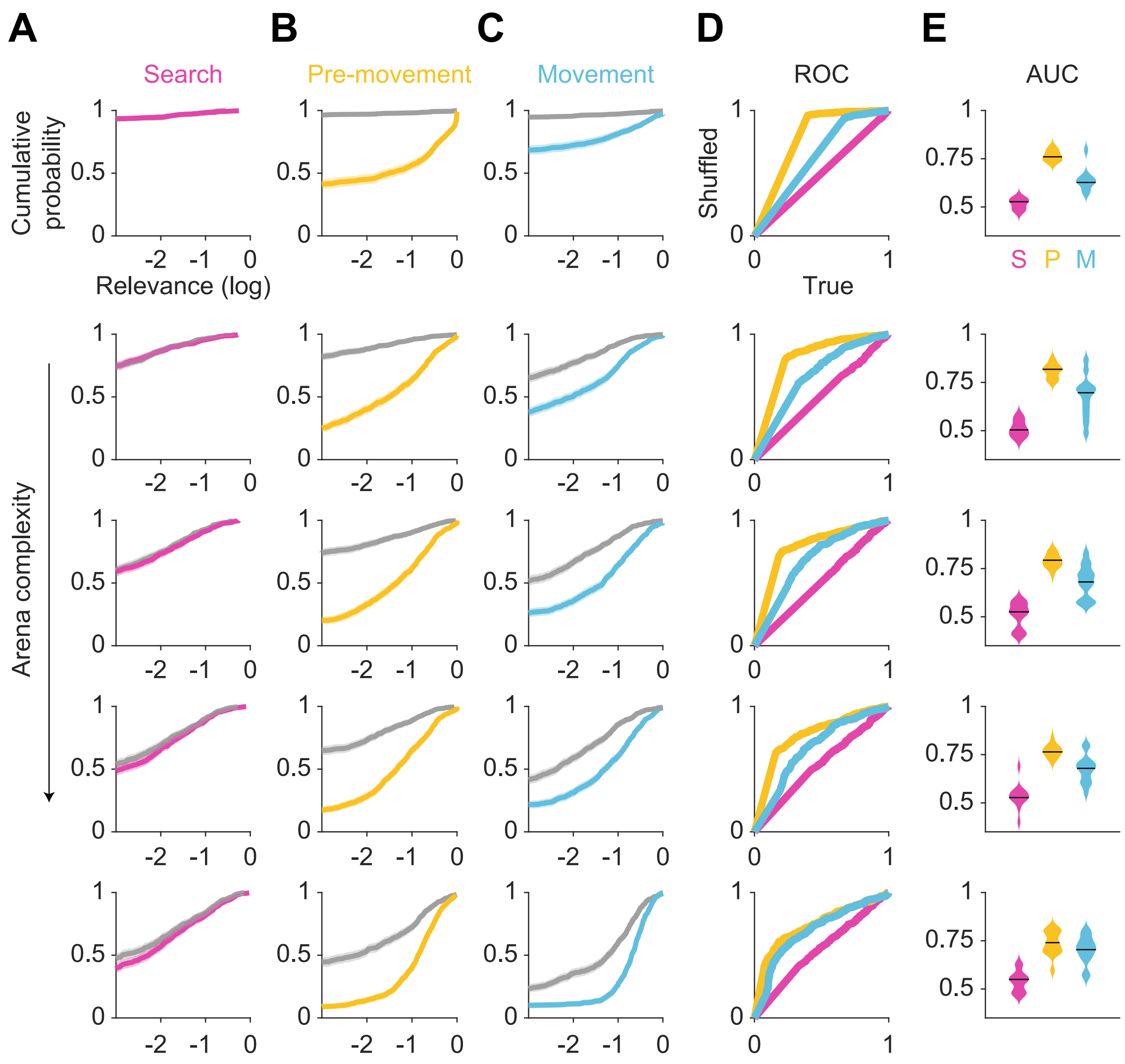


*Figure S7:* **A-E:** Breakdown of Figures 3a, b, and e for arenas 1 (most complex) through 5 (least complex).

970 **Video S1: Six representative trials in which participants exhibited sweeping eye movements.** (Top) Aerial view of the arena with the participant's dynamically evolving position (lilac) and gaze (green). The target is represented as a black circle. (Bottom) Time-evolving version of the plot described in Figure 4b. The video speed is veridical, and the search epoch was omitted from each trial.

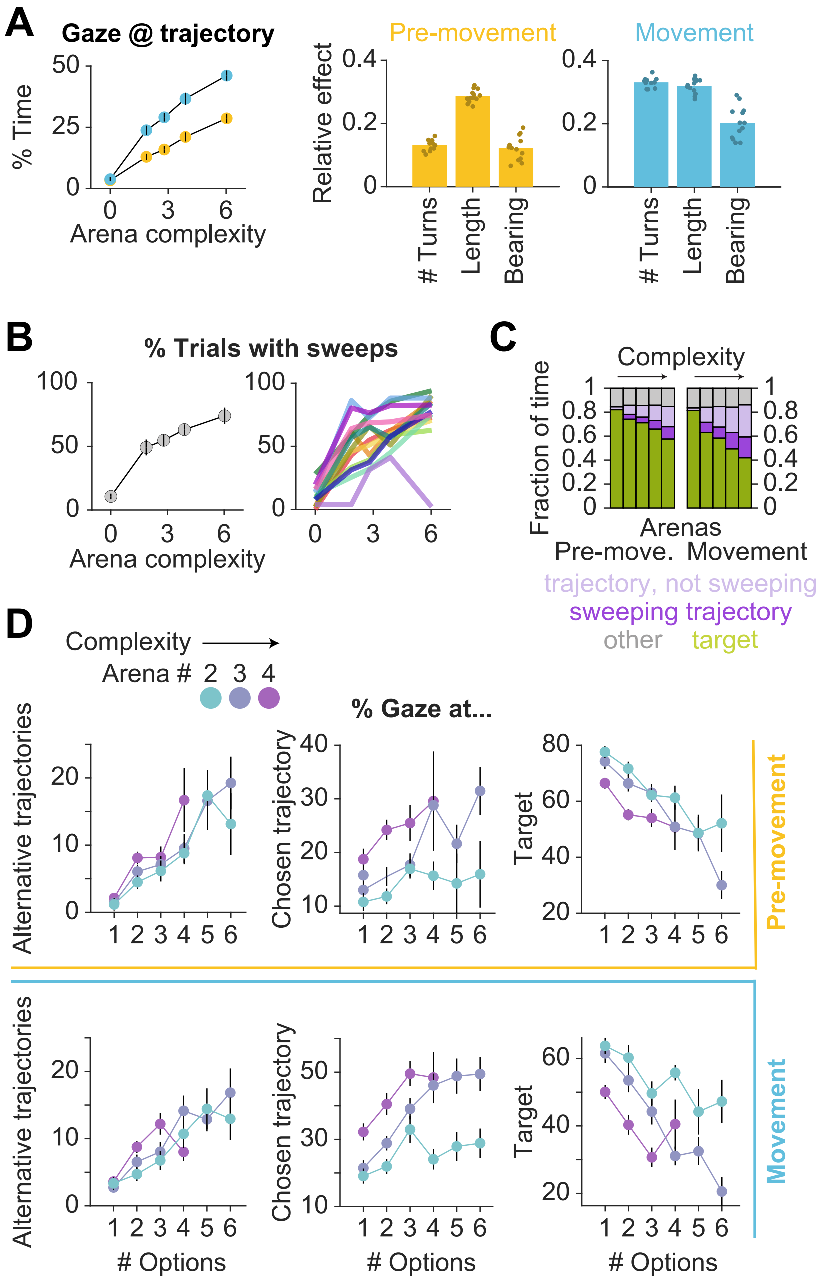


*Figure S8:* **A.** Line plot: The fraction of time that participants spent gazing near (within 2 m from) the trajectory that they took on each trial (excluding points of gaze near the goal location) increased with arena complexity during both pre-movement (gold) and movement (blue). Bar graphs: Linear mixed models with random intercepts and slopes for the effect of trial-specific variables on the fraction of time participants gazed upon the trajectory. (Please see the description for Figure S1f for further details.) The tendency to make on-trajectory eye movements increased with trial difficulty, and the expected trajectory length had the greatest effect on this statistic prior to movement. **B.** Left: The fraction of trials with sweeps was lower for less complex arenas. Right: Each color corresponds to a single participant. **C.** Eye movements on each trial were decomposed into fixations near the target (green), fixations near the participant's trajectory (excluding the target) during sweeps vs. outside of sweeps, and fixations outside of sweeps that were neither made to the target nor trajectory. Participants viewed the hidden target location more in easier arenas, and gazed upon the rest of the trajectory more in difficult arenas. The fraction of time that was spent looking elsewhere was relatively constant across arenas and epochs. **D.** During pre-movement (top row) and movement (bottom row), participants spent more time foveating alternative trajectories on trials where there were more trajectory options (left), as well as more time foveating the chosen trajectory (middle). Participants spent less time foveating the goal location when there were more trajectory options (right). All error bars denote $\pm 1$ SEM.

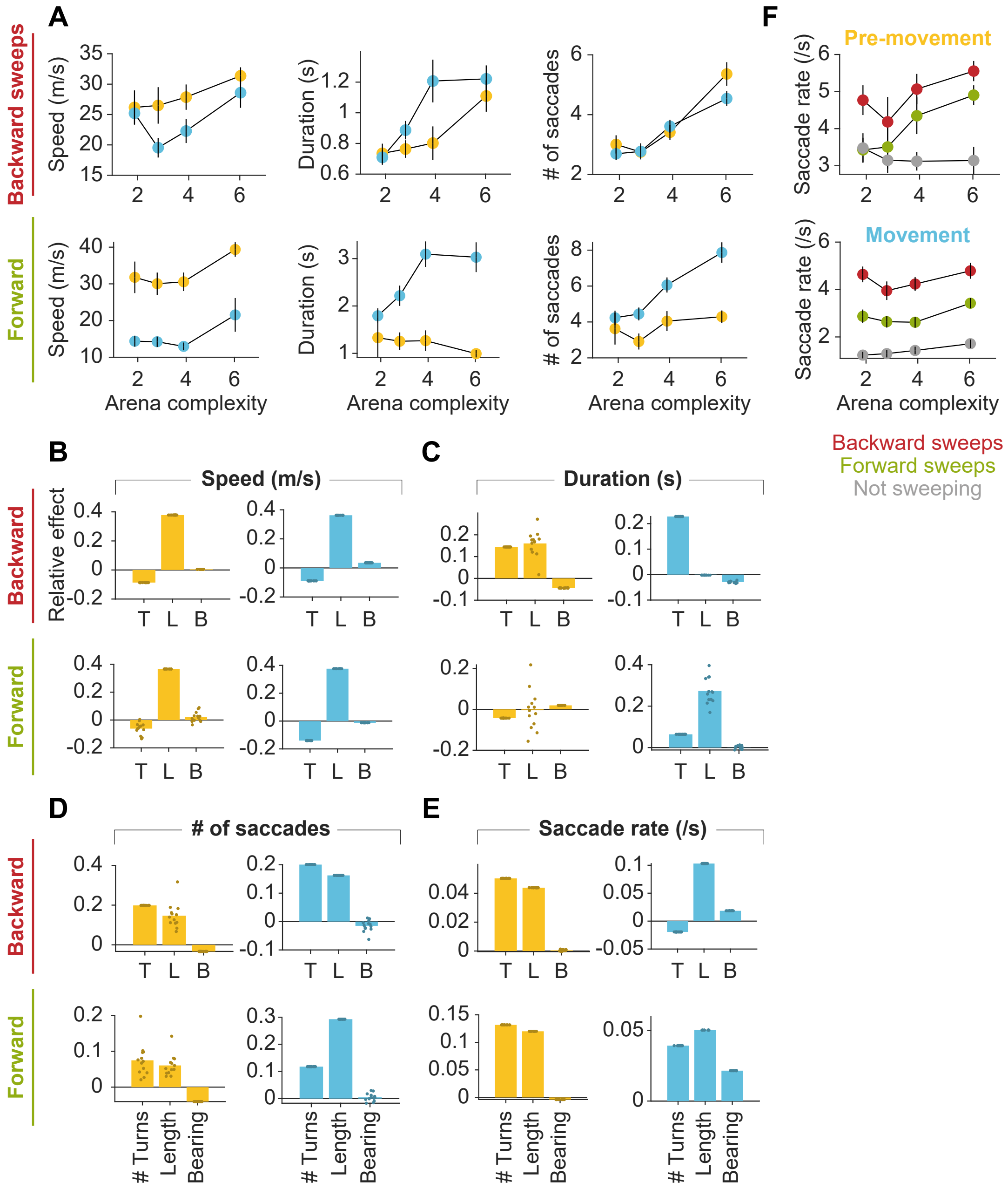


*Figure S9:* **A.** Left: Across all participants and all trials, the speed of backward sweeps (top plot) and forward sweeps (bottom plot) during pre-movement (gold) was greater than that during movement (blue). Center: Forward sweeps before movement were relatively constant in duration across different arenas (bottom plot). Otherwise, sweeps were generally longer in duration in more complex arenas, and forward sweeps were longer than backward sweeps (top plot). Right: The number of saccades during forward sweeps before movement were relatively constant across arenas. Otherwise, the average number of saccades per sweep was higher for more complex arenas. **B-E:** Linear mixed models with random intercepts and slopes for the effect of trial-specific variables on the dependent variables: **B.** speed of sweeps (m/s), **C.** duration of sweeps (s), **D.** number of saccades during each sweep, and **E.** saccade rate during sweeps. (Please see the description for Figure S1f for further details.) All variables were z-scored prior to model fitting. **F.** During both pre-movement (top plot) and movement (bottom), the saccade rate was generally higher during sweeps than outside of sweeps.

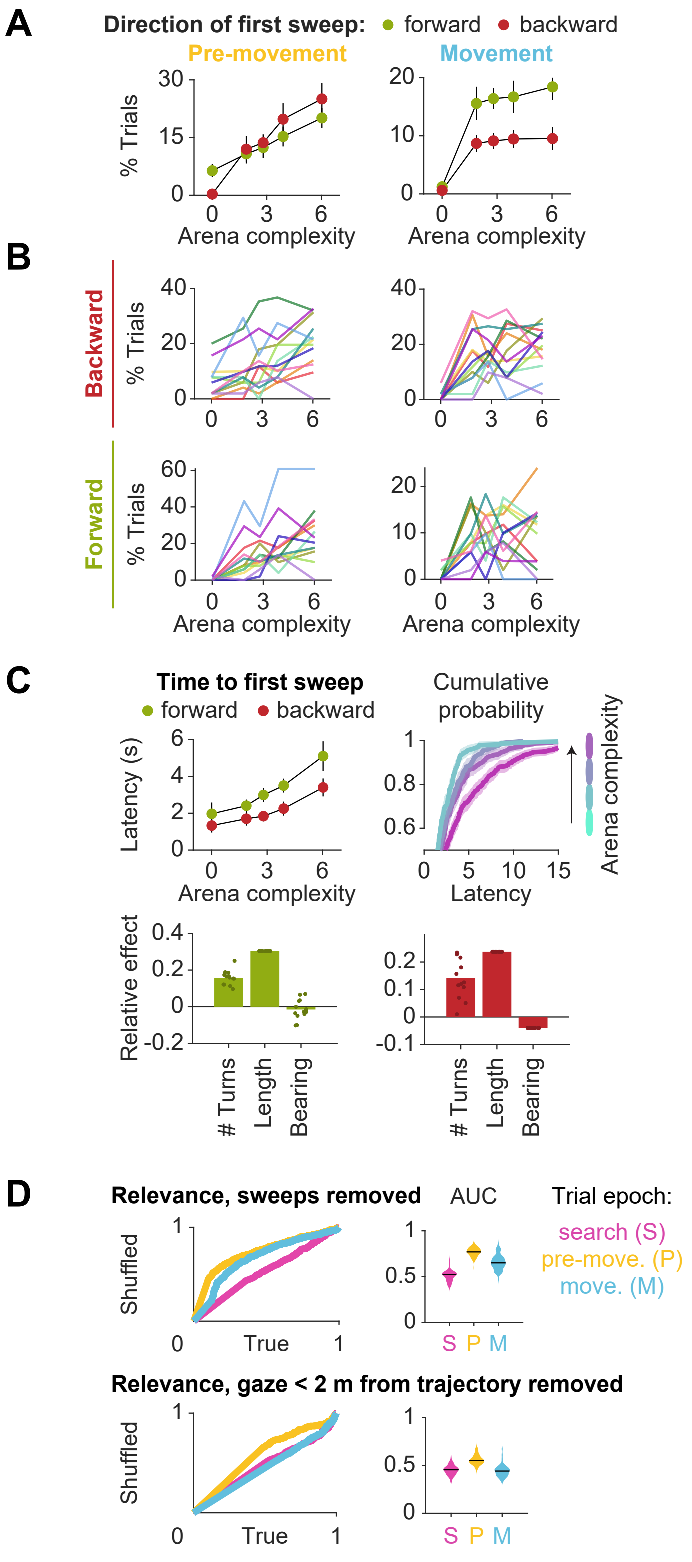

A
Direction of first sweep: forward backward
Pre-movement
Movement
% Trials
Arena complexity
B
Backward
Forward
% Trials
Arena complexity
C
Time to first sweep
forward backward
Latency (s)
Arena complexity
Cumulative probability
Latency
Arena complexity
Relative effect
### Turns
Length
Bearing
D
Relevance, sweeps removed
AUC
Trial epoch:
search (S)
pre-move. (P)
move. (M)
Shuffled
True
S P M
Relevance, gaze < 2 m from trajectory removed
Shuffled
True
S P M

*Figure S10:* **A.** The direction of the first sweep was more likely to be backwards if it occurred prior to movement (left plot), and forwards if it occurred during movement (right). **B.** Same plots as in **A**, but backward and forward sweeps are shown separately, and each color corresponds to a single participant. **C.** Top left: The average delay between goal detection and the first sweep increased with arena difficulty. Top right: Cumulative distribution of sweep delays across all participants and all trials. Bar graphs: Linear mixed models with random intercepts and slopes for the effect of trial-specific variables on the latency to the first sweep if it occurred in the forward direction (bottom left) or the backward direction (bottom right). (Please see the description for Figure S1f for additional information.) The number of turns and especially the path length positively influence sweep latency. **D.** Top left: ROC curves constructed as described in Figure 3a (rightmost) for the distributions of true vs. shuffled average relevance values for each trial (pooled across all arenas, all participants, and all trials), with periods of sweeping eye movements removed, reveals that during the pre-movement (orange) and movement (blue) epochs, non-sequential eye movements are still directed towards task-relevant locations. Top right: AUC plots were constructed with sweeps removed, as described in Figure 3b. The AUC values remain well above chance during the pre-movement (orange) and movement (blue) epochs. Bottom left/right: The same analysis was performed with gaze positions falling within 2 m of the participant's trajectory on each trial removed, revealing that the remaining visual samples were made to relevant locations in space during pre-movement, but not during movement.

---

| **Participant ID** | Block 1 | Block 2 | Block 3 | Block 4 | Block 5 |
|---|---|---|---|---|---|
| **1** | 3 | 2 | 1 | 4 | 5 |
| **2** | 5 | 4 | 1 | 2 | 3 |
| **3** | 5 | 3 | 1 | 2 | 4 |
| **4** | 4 | 5 | 2 | 3 | 1 |
| **5** | 3 | 4 | 2 | 5 | 1 |
| **6** | 5 | 2 | 1 | 3 | 4 |
| **7** | 3 | 2 | 5 | 1 | 4 |
| **8** | 4 | 5 | 3 | 1 | 2 |
| **9** | 2 | 1 | 4 | 5 | 3 |
| **10** | 5 | 3 | 1 | 2 | 4 |
| **11** | 2 | 3 | 4 | 5 | 1 |
| **12** | 1 | 2 | 5 | 3 | 4 |
| **13** | 3 | 2 | 5 | 1 | 4 |

*Table S1:* The order of arena presentation was randomized across participants.

| **Epoch** | Arena 1 | Arena 2 | Arena 3 | Arena 4 | Arena 5 |
|---|---|---|---|---|---|
| **search** | 0 (vs. 0) | 0 (vs. 0) | 0 (vs. 0) | 1.4 (vs. 0.1) | 5.3 (vs. 1.4) |
| **pre-movement** | 32 (vs. 0) | 31 (vs. 0) | 45 (vs. 0) | 47 (vs. 0) | 137 (vs. 6.2) |
| **movement** | 0 (vs. 0) | 9.5 (vs. 0) | 47 (vs. 0.7) | 51 (vs. 4.3) | 201 (vs. 60) |

*Table S2:* Median true relevance values (vs. median shuffled relevance) for each arena (x$10^{-3}$). Arenas 1-5 are in the order of least to greatest complexity.

| Figure(s) | Dependent variable | # Turns | Length | Bearing | # Options |
|---|---|---|---|---|---|
| **1i** | Pre-movement epoch duration | 12 | 12 | 10 | 1 |
| **1i** | Movement epoch duration | 13 | 13 | 12 | 5 |
| **2b** | Variance of gaze pre-move. | 13 | 13 | 11 | 5 |
| **2b** | Variance of gaze move. | 10 | 13 | 7 | 6 |
| **2d** | Gaze @ goal duration pre-move. | 13 | 13 | 10 | 8 |
| **2d** | Gaze @ goal duration move. | 13 | 13 | 12 | 10 |
| **2f** | Gaze distance from goal pre-move. | 13 | 13 | 11 | 7 |
| **2f** | Gaze distance from goal move. | 13 | 13 | 11 | 7 |
| **4c, 4d** | % Time sweeping backward pre-move. | 10 | 8 | 7 | 2 |
| **4c, 4d** | % Time sweeping backward move. | 12 | 12 | 4 | 3 |
| **4c, 4e** | % Time sweeping forward pre-move. | 2 | 3 | 3 | 2 |
| **4c, 4e** | % Time sweeping forward move. | 13 | 12 | 12 | 4 |

*Table S3:* Number of participants (13 total) with a significant Pearson's correlation ($p \leq 0.05$) between the dependent variable and each independent variable (Number of turns, Path length, Bearing angle, Number of trajectory options). Note: This correlation analyses does not characterize conditional dependencies, which may also be present in the data. Such dependencies are factored into the LME model described elsewhere.

| Figure(s) | Dependent variable | Mean slope | CV slope |
|---|---|---|---|
| **2d** | Gaze @ goal duration move. by # turns remaining | 0.80 | 5.6e-15 |
| **2f** | Gaze distance from goal move. by # turns remaining | -0.86 | -2.2e-3 |

*Table S4:* Number of participants (13 total) with a significant Pearson's correlation ($p \leq 0.05$) between the dependent variable and arena complexity (for analyses which require pooling trials). A linear mixed effects model (LME) with random slopes and intercepts yielded participant-specific slopes, from which we computed the mean and coefficient of variation (CV). Note that results showed low between-participant variability. All variables were z-scored prior to model fitting.

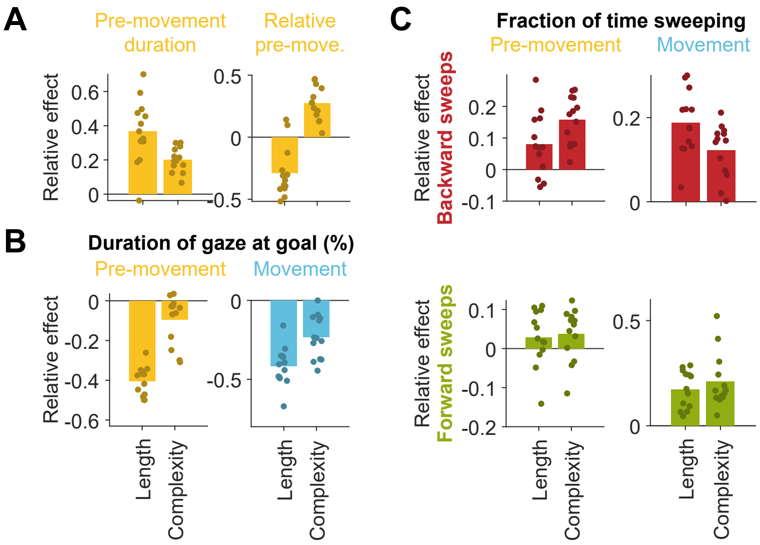


*Figure S11:* **A.** Slopes obtained by linear regression of the pre-movement epoch duration (left) or relative pre-movement epoch duration (right) against path length and arena complexity, averaged across participants. Overlaid scatter depicts participant-specific slopes. Both arena-level and trial-level variables influence these dependent variables, and notably exert opposing effects in the case of relative pre-movement duration. **B.** Left: The negative trend in the duration of gaze at the goal vs. arena complexity during pre-movement observed in Figure 2d can be mostly explained by the longer trajectories in more complex arenas. Right: However, the stronger negative trend during movement can be predicted by both arena complexity and path lengths. **C.** The fraction of time sweeping in the backwards direction pre-movement can be more strongly explained by arena-level effects than trial path lengths (top left). Both arena complexity and path lengths influence the overall fraction of time sweeping during movement (right) and the fraction of time sweeping forward pre-movement (bottom left).

**Relevance derivation:** In this section, we derive a general measure to quantify the relevance of transitions with respect to the task of navigating between two given states. The following derivation focuses on the general setting when external noise (stochastic transitions) is present and internal noise (model uncertainty) is inhomogeneous. As we will show, the measure used to quantify transition relevance in the main text (Equation 1) corresponds to the special case where transitions are deterministic and uncertainty is homogeneous. Let $T_k$ denote the status of the $k^{th}$ stochastic transition (1 or 0) and $p_k$ be the parameter of the true probability distribution (p.d.) of that transition such that $P(T_k = 1) = p_k$ and $P(T_k = 0) = 1 - p_k$. Let $\hat{p}_k$ be the parameter of the subjective probability distribution of the transition. I.e. The agent thinks that $P(T_k = 1) = \hat{p}_k$ and $P(T_k = 0) = 1 - \hat{p}_k$. Given a particular goal state, let $V_s^k$ denote the value of the agent's current state $s$ evaluated using the true transition status $T_k$ such that $V_s^k = V_s(T_k = 1)$ if $T_k = 1$ and $V_s^k = V_s(T_k = 0)$ if $T_k = 0$. Let $\hat{V}_s^k$ denote the expectation of the value of state $s$ evaluated using the subjective transition p.d. of the $k^{th}$ transition such that $\hat{V}_s^k = \hat{p}_k \, V_s(T_k = 1) + (1 - \hat{p}_k) \, V_s(T_k = 0)$. Since looking at a transition will dramatically reduce the uncertainty about the status of that transition, this can impact the subjective value of the current state, provided that transition is critical to the task at hand. For instance, discovering a subway line linking your neighborhood and downtown will increase the value of your neighborhood if your workplace is located downtown, but will have no impact if your workplace is located crosstown. Therefore, we define relevance ($\Omega_k$) of the $k^{th}$ transition as the expectation of the (log) change in subjective value about the current state $s$ induced by looking at that transition. Then, we have:

$$
\begin{aligned}
\Omega_k &= \mathbb{E}[\,\text{log}(|V_s^k - \hat{V}_s^k|)\,]_{p_k} \text{ where } \mathbb{E}[\,.\,]_{p_k} \text{ denotes expectation taken w.r.t the true p.d.} \\
&= \mathbb{E}[\,\text{log}(|V_s^k - \hat{p}_k \, V_s(T_k = 1) - (1 - \hat{p}_k) \, V_s(T_k = 0)|)\,] \\
&= p_k \, \text{log}(|V_s(T_k = 1) - \hat{p}_k \, V_s(T_k = 1) - (1 - \hat{p}_k) \, V_s(T_k = 0)|) + \\
&\qquad (1 - p_k) \, \text{log}(|V_s(T_k = 0) - \hat{p}_k \, V_s(T_k = 1) - (1 - \hat{p}_k) \, V_s(T_k = 0)|) \\
&= p_k \, \text{log}((1 - \hat{p}_k)|\Delta V|) \; + \; (1 - p_k) \, \text{log}(\hat{p}_k |\Delta V|) \text{ where } \Delta V = V_s(T_k = 1) - V_s(T_k = 0) \\
&= p_k \, \text{log}(1 - \hat{p}_k) \; + \; (1 - p_k) \, \text{log}(\hat{p}_k) \; + \; \text{log}(|\Delta V|) \\
&= (p_k - 1 + 1) \, \text{log}(1 - \hat{p}_k) \; - \; p_k \, \text{log}(\hat{p}_k) \; + \text{log}(\hat{p}_k) \; + \; \text{log}(|\Delta V|) \\
&= -(1 - p_k) \, \text{log}(1 - \hat{p}_k) \; - \; p_k \, \text{log}(\hat{p}_k) \; + \text{log}(\hat{p}_k) \; + \text{log}(1 - \hat{p}_k) \; + \; \text{log}(|\Delta V|) \\
&= \text{H}(p_k, \hat{p}_k) \; + \; \text{log}(\hat{p}_k \, (1 - \hat{p}_k)) \; + \; \text{log}(|\Delta V|) \\
&\qquad \text{where H}(X, Y) \text{ denotes the cross entropy between } X \text{ and } Y \\
&= \text{H}(p_k) \; + \; D_{\text{KL}}(p_k || \hat{p}_k) \; + \; \text{log}(\hat{p}_k \, (1 - \hat{p}_k)) \; + \; \text{log}(|\Delta V|) \\
&\qquad \text{where H}(X) \text{ denotes the entropy of } X \\
&= \text{H}(p_k) \; + \; D_{\text{KL}}(p_k || \hat{p}_k) \; + \; \text{log}(\text{Var}[\hat{T}_k]) \; + \; \tfrac{1}{2}\text{log}(|\Delta V|^2) \\
&\qquad \text{where } \hat{T}_k \text{ denotes the subjective knowledge about the status of the } k^{th} \text{ transition}
\end{aligned}
$$

Observe that $\Omega_k$ is comprised of four factors: (**I**) H$(p_k)$, the entropy of $p_k$, which captures transition volatility, (**II**) $D_{\text{KL}}(p_k || \hat{p}_k)$, the Kullback-Leibler divergence between true and subjective p.d., which captures the degree of mismatch between the subjective and true transition models, (**III**) log(Var$[\hat{T}_k]$), the log variance of the subjective status of the transition, which captures the agent's uncertainty, and (**IV**) $\text{log}(|\Delta V|^2) = \text{log}([V_s(T_k = 1) - V_s(T_k = 0)]^2)$, the log change in the value of the current state induced by changing the transition status, which captures the sensitivity of the value function to the transition. The first and third terms suggest that an agent should prioritize looking at transitions with high volatility and high subjective uncertainty. The second term suggests that it is best to look at transitions whose subjective status is known to be wrong. Although this is mathematically correct, agents would not know the true model to begin with and therefore cannot direct their attention at such transitions. Therefore, if external and internal noise are homogeneous, the best strategy would be to look at transitions which the value function is highly sensitive to, as postulated by Equation 1 in the main text. Note that in deriving $\Omega_k$, we have neglected the contribution of model mismatches that may exist at other transitions. This approximation will be valid if the model mismatch is small, and the solution works well in practice as demonstrated by the simulations (Figure S5).
